# Ventral Hippocampus and NAc Core, but not Shell activity is required for well-learned instrumental memory destabilization

**DOI:** 10.1101/2025.05.21.654887

**Authors:** Ramon Hypolito lima, Chaoran Cheng, Jonathan L. C. Lee

## Abstract

While there is increasing evidence that instrumental memories undergo destabilization and reconsolidation, little is known about the underlying neural mechanisms of these processes. Here, we focussed on instrumental memory destabilization, and the functional involvement therein of the nucleus accumbens core and shell regions, as well as the ventral hippocampus. In male rats, and using a previously established destabilization reminder procedure, we infused the GABA receptor agonists baclofen/muscimol at the time of memory reminder. We found infusions into the core but not shell impaired well-learned instrumental memory destabilization, protecting against MK-801-induced reconsolidation impairment. Moreover, infusion of baclofen/muscimol into the ventral hippocampus resulted in the same protective effect. These results suggest that the NAc core, but not the shell or vHPC, is involved in the destabilization of well-learned instrumental memory, potentially through a vHPC-NAc core projection.

## Introduction

The formation and stabilisation of a stable mnemonic trace depends on several important stages, including acquisition, encoding, consolidation, storage, and potential retrieval and behavioural expression. Within this framework, the hypothesised process of memory reconsolidation is argued to underpin the modification of previously consolidated memories (Lewis, 1979; Nader, 2003), but see (Schroyens, Beckers and Luyten, 2022). Memory reconsolidation involves the destabilisation of an established long-term memory following a reminder (Milton, Das and Merlo, 2023). When destabilised, the memory is rendered temporarily malleable, and its updated form may be restabilised into long-term memory (Lee, 2009).

For instrumental memories, there is growing evidence that memory reminder can trigger memory destabilisation, such that the administration of NMDA receptor antagonists results in subsequent impaired instrumental performance (Exton-McGuinness *et al*., 2014, Exton-McGuinness *et al*., 2019; Tedesco *et al*., 2014; Exton-McGuinness and Lee, 2015; Cheng, Exton-McGuinness and Lee, 2022). Typically, these studies have used systemic administration of MK-801 as the amnestic agent, although the nature of memory reminder has differed between studies (Piva *et al*., 2020). While simple extinction training has been shown to be sufficient under some circumstances (Tedesco *et al*., 2014), the use of an altered instrumental contingency at memory reminder may be a more reliable means of triggering memory destabilisation (Exton-McGuinness *et al*., 2014; Exton-McGuinness and Lee, 2015).

The systemic route of MK-801 administration gives little insight into the neural substrates of instrumental memory destabilisation and reconsolidation. Previous immunohistochemical analyses have shown an upregulation of Zif268 expression in the nucleus accumbens (NAc), as well as the amygdala (Piva *et al*., 2019). While these regions are consistently implicated in instrumental memory consolidation and performance (Kelley, Smith-Roe and Holahan, 1997; Baldwin *et al*., 2000; Smith-Roe and Kelley, 2000; Balleine, Killcross and Dickinson, 2003; Dalley *et al*., 2005), and have also been shown to be critical loci of memory destabilisation and reconsolidation for non-instrumental memories (Miller and Marshall, 2005; Ben Mamou, Gamache and Nader, 2006; Ren *et al*., 2013; Liang *et al*., 2017), it remains difficult to draw strong conclusions about their necessity for instrumental memory destabilisation and/or reconsolidation. Given that instrumental learning and performance occurs in parallel with discrete and contextual pavlovian appetitive learning, memory reminder may trigger destabilisation of any or all of these instrumental and pavlovian associations (Cheng, Exton-McGuinness and Lee, 2022). Therefore, studies that show impairments in instrumental memory destabilisation/reconsolidation are necessary to provide evidence of neuroanatomical involvement.

Here we used intracranial infusions of the GABA receptor agonists baclofen and muscimol to functionally inactivate target regions. When coupled with instrumental memory reminder, via contingency change, and MK-801 injection, this approach tests whether baclofen/muscimol infusion prevents memory destabilisation to protect against MK-801-induced reconsolidation impairment. Infusions were targeted at the NAc core and shell regions, given their implication in instrumental memory consolidation and performance (Corbit, Muir and Balleine, 2001; Shiflett, Brown and Balleine, 2010; Ostlund *et al*., 2011), and the ventral hippocampus as a non-striatal region that functionally projects to the nucleus accumbens (Floresco, Todd and Grace, 2001) and is necessary for appetitive memory expression (Rogers and See, 2007; Zhou *et al*., 2020).

## Methods

### Animals

Subjects were 83 male experimentally-naive Lister Hooded rats (Charles River, UK) weighing 300-350 g at the time of surgical procedures. They were housed in a standard facility in individually ventilated cages under a normal light cycle (12 hr:12 hr, lights on at 0700). Cages contained Aspen chip bedding and a plexiglass tube, a cardboard house and wooden chew sticks for environmental enrichment. Water was available *ad libitum*; standard lab chow was restricted to 20 g/rat/day from the first day of behavioural procedures until the end of the experiment. Rats were fed immediately after the behavioural session. All procedures were approved by the local Animal Welfare and Ethical Review Board, and performed under the authority of the Animal (Scientific Procedures) Act 1986 (Amendment Regulations 2012)(PPL P3B19B9D2).

### Surgical procedures

Under aseptic conditions, rats were implanted with bilateral chronic indwelling stainless steel cannulae as previously described (Exton-McGuinness and Lee, 2015). Bilateral cannulae targeted the NAc core (AP +2.0 mm, ML ±1.5 mm, DV −3.0 mm), NAc shell (AP +2.2 mm, ML ±0.9 mm, DV −3.0 mm) or vHPC (AP −5.3 mm, ML ±4.5 mm, DV −4.0 mm); all DV coordinates relative to skull surface. Rats received perioperative analgesia through subcutaneous injection of buprenorphine and free access to carprofen gel. Rats were housed individually in temperature-controlled cages for the first night after cannula implantation and then were returned to their group-housed cage. Behavioural procedures began 8-9 days after surgery, during which time the rats were weighed and handled regularly.

### Drugs

MK-801 (AbCam, UK) was dissolved in sterile saline to a concentration of 0.1 mg/mL. Rats were injected intraperitoneally (i.p.) with 0.1 mg/kg of MK-801 or an equivalent volume of saline vehicle, 30 min prior to the reminder session (Nac infusion experiments) or immediately after the reminder session (vHPC infusion experiment).

For intracerebral infusions, rats received a combined Baclofen/Muscimol solution (0.1 mM muscimol/1.0 mM baclofen; 0.5 μl/side, 0.25 μl/min) or sterile PBS vehicle (0.5 μl/side, 0.25 μl/min) immediately before the reminder session. Injectors projected beyond the guide cannulae by 2.0 mm (vHPC), 3.4 mm (NAc core) and 3.8 mm (NAc shell).

### Behavioural procedure

The behavioural procedures were carried out in eight operant chambers (MedAssociates), each measuring 25 × 32 × 25.5 cm and individually housed within a sound-attenuating enclosure, as previously described (Cheng, Exton-McGuinness and Lee, 2022) and established as a robust method for inducing instrumental memory destabilisation (Figure 1).

**Figure 1.**
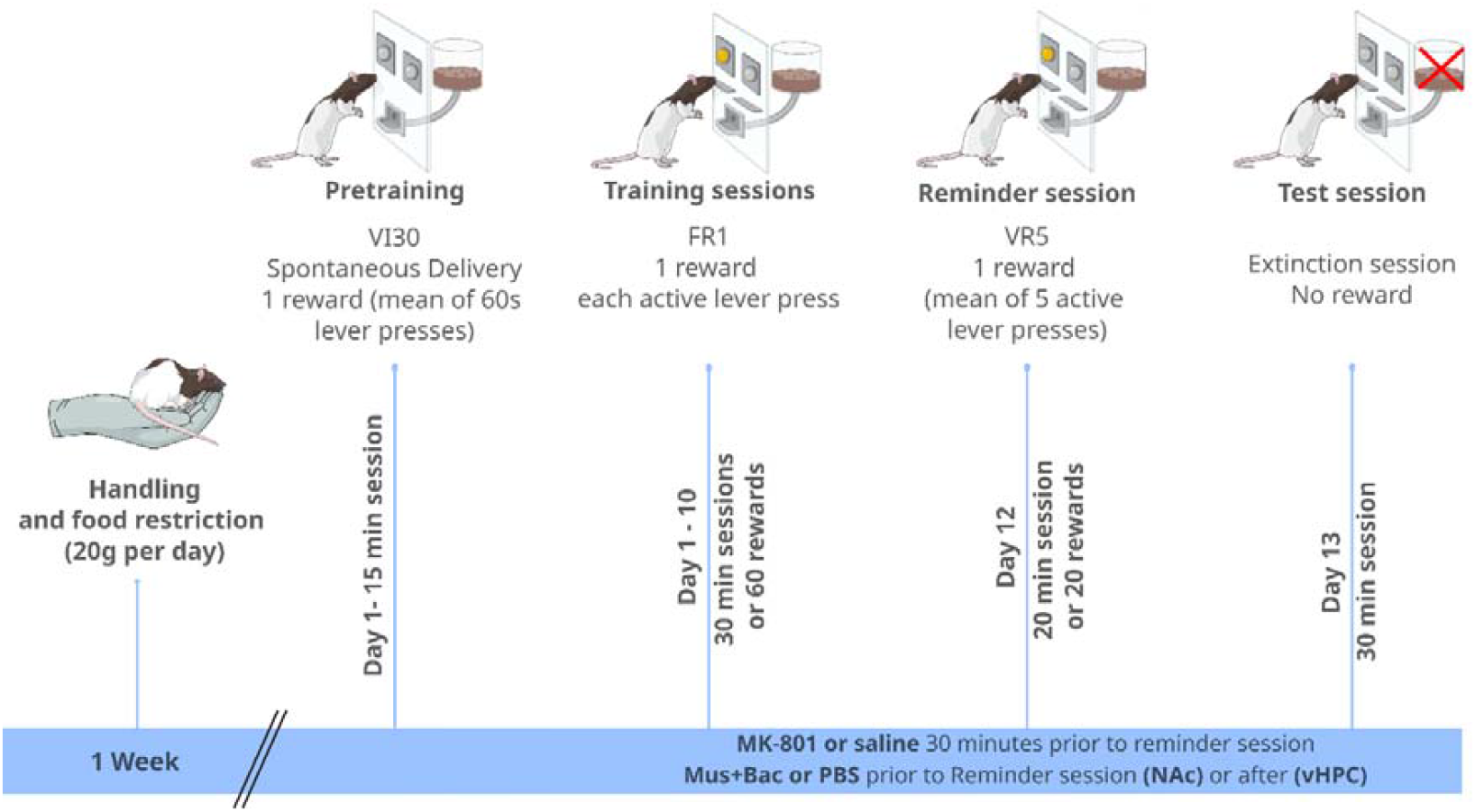
Schematic illustration of behavioural procedures. Animals were subjected to a food restriction protocol (20_g/rat/day) and daily handling sessions for one week prior to the onset of behavioral training. Rats were then trained over 10 consecutive days. On the first training day, all animals underwent a VI30 (variable interval 30_s) pre-training session immediately before the initial training session. A reminder session under a VR5 (variable ratio 5) schedule was conducted 48 hours after the final training day. Rats received intraperitoneal injections of MK-801 or saline either 30 minutes before (for NAc experiments) or immediately after (for vHPC experiments) the reminder session. Additionally, infusions of muscimol/baclofen (Mus/Bac) or PBS were administered prior to the reminder session. The test session was conducted 24 hours following the reminder session.

### Training Sessions

Rats were exposed to a variable-interval (VI) schedule pretraining session (mean, 60 sec; range, 30–90 sec), during which animals were allowed to collect 45-mg sucrose rodent pellets (5TUL, TestDiet) from the magazine for 15 min. This session was used to facilitate instrumental learning over the subsequent training sessions.

The first instrumental training session began immediately after the pretraining session. At the beginning of each training session, both levers were presented to the animals in the box. Active lever presses (assigned pseudorandomly) delivered a reward pellet on an FR1 schedule (one lever press delivered one pellet); responses on the lever had no other programmed consequence, the levers did not retract and remained extended throughout the session and no discrete reward-predictive stimuli were presented at any point during training. Instrumental training sessions lasted 30 min or until a maximum of 60 pellets had been obtained. Rats received a total of 10 training sessions (with a maximum of 60 rewards each) over 10 consecutive weekdays.

### Reminder session

Two days after the final training session, rats were given a VR5 reminder session. A variable number of active lever presses (mean: 5, range: 1-9) were required to obtain a reward. The session lasted for 20 minutes, or when 20 sucrose pellets were obtained – whichever occurred first. This reminder was chosen based on previous findings that variable-ratio schedules destabilise a well-established instrumental memory (Exton-McGuinness *et al*., 2014; Cheng, Exton-McGuinness and Lee, 2022). Infusions and injections were administered as described above.

### Post-reminder test

Instrumental performance was tested the day after drug treatment for all groups. Test sessions lasted 30 min and were performed in extinction. The levers were extended, no rewards were delivered, and the house light remained on throughout the session.

### Cresyl violet staining and verification of the cannulae placements

Rats were killed at the end of behavioural procedures and their brains were carefully removed and stored in 4% paraformaldehyde (PFA) solution overnight at 4^°^C. The brains were sliced at 40 um and stained with cresyl violet using a standard procedure. Brain slices were observed using an Olympus BX50 light microscope attached to a Leica DFC425 camera.

### Statistical analysis

Data are presented as mean + S.E.M. for the number of lever presses at each training, retrieval, and test sessions. All data were checked for consistency, and any data lying more than two standard deviations from the mean were treated as outliers and excluded (2 outliers was excluded in vHPC B/M-saline group). Data were analysed using repeated-measures analysis of variance (ANOVA; vHPC, B/M-MK-801 n =7, B/M-saline n= 5, PBS-MK-801 n=8, PBS-saline n = 7, Nac core, B/M-MK-801 n =7, B/M-saline n= 7, PBS-MK-801 n=7, PBS-saline n = 7, Nac shell, B/M-MK-801 n =7, B/M-saline n= 7, PBS-MK-801 n=7, PBS-saline n = 7), with Day (as appropriate), Infusion and Injection as factors. Results with p < 0.05 were deemed significant. A Greenhouse–Geisser correction was used to correct for nonspherical data (as assessed by Mauchley’s Test of Sphericity).

## Results

### core

At test, pre-reminder MK-801 resulted in reduced active lever pressing, but not in intra-NAc core B/M-infused rats (Figure 2B; Inf x MK-801 x Lever: F(1,24)=6.46, p=0.018, η^2^_p_=0.21, BF_inc_=9.3; simple main effects of MK-801 in PBS active lever only (p=0.003; all other p’s>0.13).

**Figure 2.**
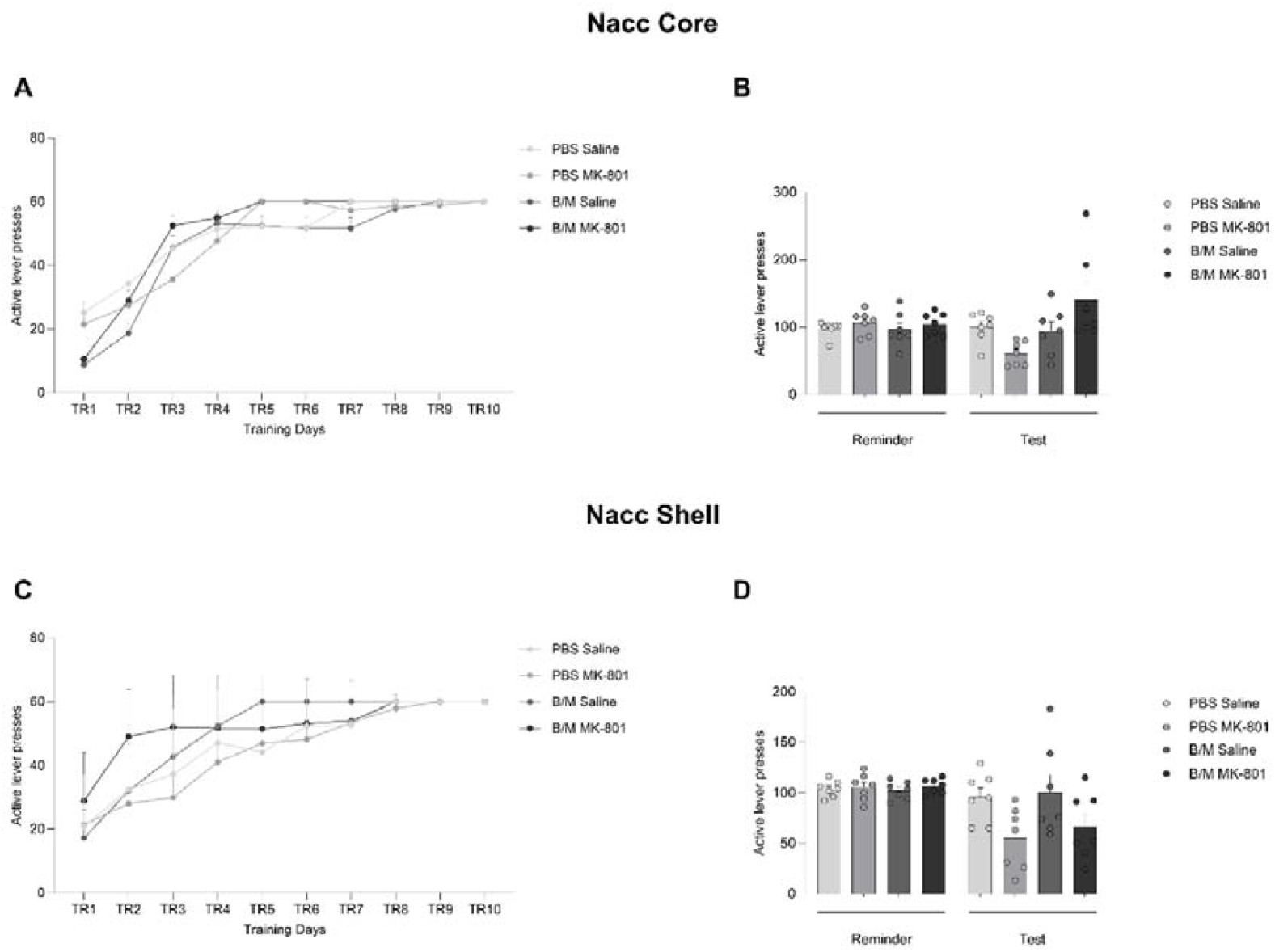
Nac Core but not Shell is required to memory destabilization. Rats were trained to press the correct lever to receive sucrose reward during 10 days, followed by a VR5 reminder session and administration of MK-801 or saline and infusion of Baclofen/Muscimol or PBS. **(A and C)** All rats learned to discriminate active and inactive lever during the training days. **(B)** pre-reminder MK-801 administration reduced active lever pressing at test, and a pre-reminder infusion of B/M intro the NAc core protected against the MK-801-induced impairment in active lever pressing at the test. **(D)** the pre-reminder infusion of B/M intro the NAc shell did not appear to protect against the MK-801-induced impairment in active lever pressing at the test. Data are presented as mean ± SEM.

This difference at test was not evident previously at the reminder session (Figure 2B; Inf x MK-801 x Lever: F(1,24)=0.11, p=0.74, η^2^_p_ =0.005, BF_inc_ =0.072; simple main effect p=0.26 for PBS active lever), nor were there any obvious differences during training that might account for the differential effects of MK-801 at test (Figure 2A ; Inf x MK-801 x Lever x Session: F(3.6,68.8)=0.36, p=0.82, η^2^_p_=0.019, BF_inc_=7.4 × 10 ^−9^; Inf x MK-801 x Lever: F(1,19)=0.38, p=0.55, η^2^_p_ =0.020, BF_inc_ =0.004). Upon inspection of the analysis only the Inf x Lever x Session interaction was marginal (F(3.6,68.8)=2.21, p=0.083, η ^2^_p_=0.10, BF_inc_=0.026) with analyses of simple main effects revealing higher lever pressing in the PBS-infused rats compared to the B/M-infused rats in the first training session only (p=0.014). However, all rats reached asymptotic levels of performance by the end of training. Therefore, the pre-reminder infusion of B/M intro the NAc core protected against the MK-801-induced impairment in active lever pressing at the test.

### NAc shell

At test, pre-reminder MK-801 resulted in reduced active lever pressing regardless of infusion into the NAc shell (Figure 2D; MK-801 x Lever: F(1,24)=7.84, p=0.010, η^2^_p_=0.25, BF_inc_=19.7; Inf x MK-801 x Lever: F(1,24)=0.10, p=0.75, η^2^_p_=0.004, BF_inc_=0.32). However, planned analyses of simple main effects revealed strong evidence for an effect of MK-801 only on active lever pressing in the PBS-infused rats (p=0.017), with weaker evidence in the B/M-infused rats (p=0.069).

At the reminder session, while there was no Inf x MK-801 x Lever interaction (Figure 2D; F(1,24)=0.00, p=1.00, η^2^_p_ =0.00, BF_inc_ =0.085), there was weak evidence for a MK-801 x Lever interaction (Figure 2D; F(1,24)=3.43, p=0.077, η^2^_p_ =0.13, BF_inc_=1.04). This appeared to be driven by the lower inactive lever pressing in rats injected with MK-801 prior to the reminder session (simple main effects p=0.014 for PBS inactive lever, all other p’s>0.27). This lower inactive lever pressing did not survive the test and so is unlikely to account for the differences observed at the test.

There were no obvious differences during training that might account for the effects of MK-801 at test (Figure 2C; Inf x MK-801 x Lever x Session: F(3.3,80.0)=1.10, p=0.36, η^2^_p_ =0.044, BF _inc_ =9.1 × 10^−10^; Inf x MK-801 x Lever: F(1,24)=0.005, p=0.94, η^2^_p_=0.00, BF_inc_=0.008). Therefore, the pre-reminder infusion of B/M intro the NAc shell did not appear to protect against the MK-801-induced impairment in active lever pressing at the test.

### vHPC

At test, post-reminder MK-801 resulted in reduced active lever pressing, but not in B/M-infused rats (Inf x MK-801 x Lever: F(1,24)=4.65, p=0.041, η^2^_p_=0.16, BF_inc_=4.0; simple main effects of MK-801 in PBS active lever only (p=0.016; all other p’s>0.31). This difference in the test was not evident previously at the reminder session (Inf x MK-801 x Lever: F(1,24)=2.84, p=0.11, η ^2^ _p_ =0.11, BF_inc_ =0.46; simple main effects p=0.87 for PBS active lever). However, there was a simple main effect of the drug injection group on active lever responding in rats that received B/M infusion. This difference did not appear to influence the response pattern at the test as the B/M-MK-801 group increased active lever responding from reminder to the test, whereas the Sal-MK-801 group substantially decreased.

There were also no obvious differences during training that might account for the differential effects of MK-801 at test (Figure 3A; Inf x MK-801 x Lever x Session: F(3.2,74.4)=1.17, p=0.33, η^2^_p_=0.048, BF_inc_=6.93 × 10^−8^; (Inf x MK-801 x Lever: F(1,23)=0.37, p=0.55, η^2^_p_ =0.016, BF_inc_ =0.012). Upon inspection of the analysis only the Inf x Lever x Session interaction was marginal (F(2.9,67.6)=2.20, p=0.097, η^2^_p_=0.087, BF_inc_=2.6 × 10^−5^) with analyses of simple main effects revealing little evidence for group differences at any of the training sessions, with all rats reached asymptotic levels of performance by the end of training. Therefore, the pre-reminder infusion of B/M into the NAc core protected against the post-reminder MK-801-induced impairment in active lever pressing at the test.

**Figure 3.**
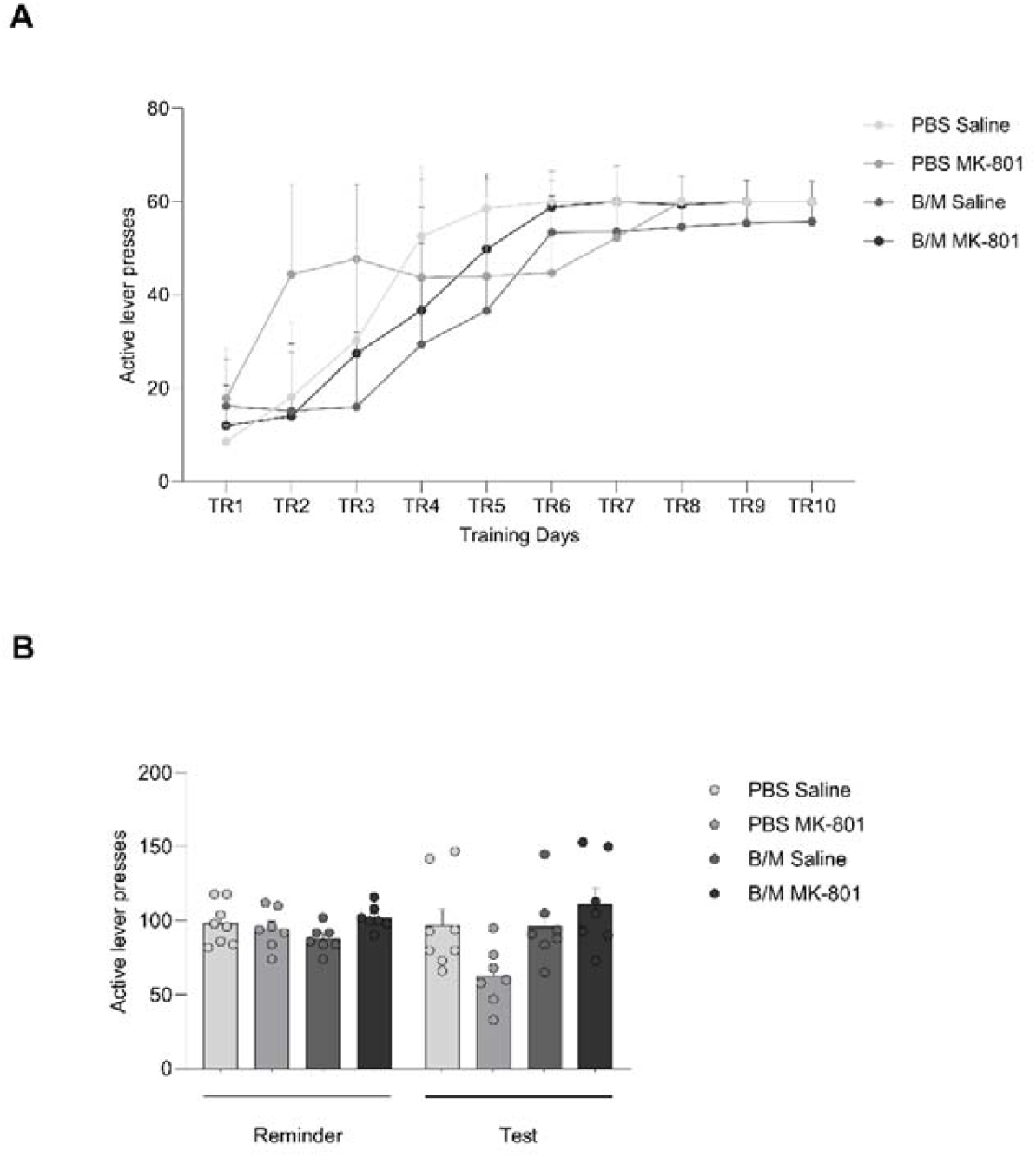
vHPC is required to memory destabilization. Rats were trained to press the correct lever to receive sucrose reward during 10 days, followed by a VR5 reminder session and administration of MK-801 or saline and infusion of Baclofen/Muscimol or PBS. **(A)** All rats learned to discriminate active and inactive lever during the training days. **(B)** post-reminder MK-801 administration reduced active lever pressing at test, and a pre-reminder infusion of B/M intro the vHPC protected against the MK-801-induced impairment in active lever pressing at the test. Data are presented as mean ± SEM.

## Discussion

Using an established procedure to impair the reconsolidation of instrumental sucrose memories, here we show that infusion of baclofen/muscimol into the NAc core or the vHPC protected against MK-801-induced impairment. In contrast, infusion into the NAc shell did not have an obvious modulatory effect on the impact of MK-801. These results suggest that neural activity in the NAc core and vHPC, but not the NAc shell, is functionally involved in the destabilisation of instrumental memories.

The use of peri-reminder injections of MK-801 to impair memory reconsolidation in instrumental reward-seeking settings is well established, especially under the reminder conditions used here. While the initial demonstration of instrumental memory reconsolidation employed a VR20 reminder session on the day after the end of single-lever instrumental training (Exton-McGuinness *et al*., 2014), our subsequent studies in a two-lever (active vs inactive) instrumental paradigm demonstrated that a VR5 reminder session two days after the end of training was the most reliable approach to destabilise the instrumental memory (Cheng, Exton-McGuinness and Lee, 2022). In our previous studies, MK-801 was injected 30 min prior to the memory reminder session (Exton-McGuinness *et al*., 2014, Exton-McGuinness *et al*., 2019; Exton-McGuinness and Lee, 2015; Cheng, Exton-McGuinness and Lee, 2022), consistent with its timing in the impairment of non-instrumental memory reconsolidation (Lee, Milton and Everitt, 2006; Brown, Lee and Sorg, 2008; Milton *et al*., 2011). MK-801 also has reconsolidation-impairing effects when administered shortly after the reminder session (Alaghband and Marshall, 2012), including in an instrumental nicotine-seeking setting (Tedesco *et al*., 2014). Here, our experiments included both pre-reminder (NAc core and shell) and post-reminder (vHPC) administration of MK-801. The observation that MK-801 impairs instrumental performance at test in the control-infused conditions further supports the conclusion that the timing of MK-801 administration is not critical in impairing memory reconsolidation, at least in our paradigm (see (Tedesco *et al*., 2014) for an example, in which pre-reminder administration is without obvious effect).

Dual-treatment approaches, in which the first treatment protects against the amnestic effect of the second treatment, are commonly interpreted as isolating the mechanisms of memory destabilisation (Ben Mamou, Gamache and Nader, 2006; Rao-Ruiz *et al*., 2011; Milton, Das and Merlo, 2023; Almeida-Corrêa and Amaral, no date). Intracranial infusions of muscimol have been used previously to impair memory destabilisation in fear (Troyner and Bertoglio, 2020) and spatial (Rossato *et al*., 2015) memory settings, and we have previously used baclofen/muscimol to block the destabilisation of a goal-tracking memory (Reichelt, Exton-McGuinness and Lee, 2013). Here, infusions of baclofen/muscimol into the NAc core, but not shell, impaired the destabilisation of the instrumental sucrose memory. While this is consistent with the pattern of selective NAc core involvement in destabilisation in a cocaine conditioned place preference paradigm (Ren *et al*., 2013), the engagement of the NAc shell following instrumental memory retrieval, and its downregulation by MK-801 administration does suggest some functional involvement of the shell in instrumental memory destabilisation and/or reconsolidation (Piva *et al*., 2018). It should be noted, however, that despite similar evidence of immediate-early gene expression in both core and shell following appetitive pavlovian memory reminder (Thomas and Everitt, 2001; Li *et al*., 2016), there remains evidence that pharmacological modulation of the core, but not the shell, impairs memory reconsolidation ((Liang *et al*., 2017; Shen *et al*., 2022), but see (Li *et al*., 2015)). Of course, these are reconsolidation impairment studies, rather than destabilisation-targeting studies, and while the neuroanatomical loci of memory reconsolidation can be targeted for destabilisation (Ben Mamou, Gamache and Nader, 2006; Merlo *et al*., 2015), the circuitry that regulates memory destabilisation appears to extend beyond the loci of underpinning plasticity (Reichelt, Exton-McGuinness and Lee, 2013; Rossato *et al*., 2015).

The functional involvement of non-mnemonic neural loci in memory destabilisation may explain our observation that intra-vHPC baclofen/muscimol also impaired instrumental memory destabilisation. The vHPC is somewhat implicated in instrumental performance. Neonatal lesions to the vHPC appear to enhance instrumental acquisition ((Brady *et al*., 2008), but see (Macedo *et al*., 2008)), and adult lesions resulted in disinhibited instrumental responding (Gourley *et al*., 2010). Moreover, the vHPC appears to be necessary for the encoding and expression of context-instrumental relations (Piquet, Faugère and Parkes, 2023), indicating an important role in the regulation of instrumental memory and performance. While it is possible that our simple, non-context-dependent, instrumental paradigm still engages context-instrumental memory processes in the vHPC, it is unlikely that this accounts directly for the patterns of behaviour in the current study, notwithstanding prior demonstrations of reconsolidation within vHPC in non-instrumental memories (Chia and Otto, 2013; Vafaei *et al*., 2023). Even were peri-reminder MK-801 injection to impair the context-instrumental association, this would not be expected to result in a reduction in instrumental performance at test. Rather, it is perhaps more likely that the change in instrumental contingency at memory reminder might engage the vHPC in a manner that regulates plasticity downstream in the NAc core. vHPC lesions or temporally-specific inhibition impaired the normal behavioural consequences of instrumental contingency degradation, indicating an important involvement in contingency monitoring and change detection (Piquet, Faugère and Parkes, 2024). Given the functional role of plasticity in the vHPC-NAc core projection in cocaine conditioned place preference retrieval and expression (Zhou *et al*., 2019), and a similar requirement for the vHPC-NAc projection (albeit not isolated to the NAc core) in reconsolidation of the cocaine place preference memory (Caban Rivera *et al*., 2023), it is plausible that a similar projection underpins the integration of the vHPC and NAc core in instrumental memory destabilisation.

In summary, here we present evidence that the NAc core and vHPC, but not the NAc shell, are functionally involved in a neural circuit that coordinates the destabilisation of instrumental sucrose memories upon a change in instrumental contingency. It remains to be determined whether the anatomical projection from the vHPC to the NAc is functionally necessary for destabilisation, as well as whether either or both areas have any role to play in the plasticity taking place during instrumental sucrose memory reconsolidation. Nevertheless, the current findings show the validity of initially focussing on the neural substrates of memory destabilisation, in the absence of any direct evidence for the neural substrates of memory reconsolidation for the memory/paradigm in question.

## Data availability statement

The raw data will be made available by the authors upon request.

## Author contributions

RHL and CC conducted the experiments. RHL, CC and JL designed the experiments, analysed the data and wrote the manuscript. All authors approved the submitted version.

## Acknowledgements

We thank David Barber for technical assistance.

